# Dopaminergic Modulation of Working Memory and Cognitive Flexibility in a Zebrafish Model of Aging-Related Cognitive Decline

**DOI:** 10.1101/2020.06.05.136077

**Authors:** Madeleine Cleal, Barbara D. Fontana, Molly Double, Roxana Mezabrovschi, Leah Parcell, Edward Redhead, Matthew O. Parker

## Abstract

Healthy aging is associated with a decline in memory and executive function, which have both been linked with aberrant dopaminergic signalling. We examined the relationship between cognitive performance and dopamine function of young and aging zebrafish (*Danio rerio*). We revealed age-related decreases in working memory and cognitive flexibility in the Free-Movement Pattern (FMP) Y-maze. An increase in *drd5* gene expression in aging adults coincided with a decrease in cognitive performance. Treatment with a D1/D5 receptor agonist (SKF-38393, 35 μM) 30 minutes prior to behavioural assessment resulted in improved working memory in aging zebrafish, but no effect in younger adults. However, an ‘overdosing’ effect caused by agonist treatment resulted in downregulation of *dat* expression in 6-month old, treated zebrafish. The translational relevance of these findings was tested in humans by analysing exploratory behaviour in young-adult, 18-35-year olds, and aged adults, 70+ year olds, in a virtual FMP Y-maze. Our findings revealed similar age-related decline in working memory. Thus, strongly supporting zebrafish as a translational model of aging and cognitive decline.

## Introduction

During the process of ‘natural’ aging, the brain undergoes gradual structural and functional changes that cause deterioration in cognitive ability, even in the absence of neurodegenerative disease (Harada et al., 2013; Salthouse, 2009). The rate of decline is highly variable, with many able to maintain good health and mental ability into their late 80s, whilst others are greatly susceptible to debilitating cognitive impairment and disease (Deary et al., 2012). Individual differences in cognitive aging have driven the identification of underlying biological factors that can predict vulnerability to decline or development of age-related diseases (Berry et al., 2016). Many studies of healthy aging have implicated multiple components of the dopaminergic system in declining cognition, with individual differences in dopamine signalling playing a profound role in performance of cognitive tasks and response to dopamine-altering medications (Roshan Cools & D’Esposito, 2011; Kimberg et al., 1997; Volkow et al., 1998). As a result, dopamine has become a focus of research investigating age-associated changes in cognition.

Dopamine plays an important role in many aspects of cerebral functions related to cognition such as attention, learning, working memory and mental flexibility (Girault & Greengard, 2004; Naderi et al., 2016). Three key modulators of dopamine function are dopamine synthesis, reuptake and activation of dopaminergic receptors (Klanker et al., 2013). Through pharmacological manipulations, a complex network of interactions involving these modulatory systems are used to maintain dopamine homeostasis (Roshan Cools & D’Esposito, 2011), dysregulation of which has been shown to have severe, and sometimes opposing, effects on cognition, particularly working memory and cognitive flexibility in health and disease (Brozoski et al., 1979; Cai & Arnsten, 1997; El-Ghundi et al., 2007; Rothmond et al., 2012; Zahrt et al., 1997). In a previous study, we sought to delineate the role dopaminergic receptor subtypes play in formulating search strategies in zebrafish exploring the FMP Y-maze. We used SCH-23390, a selective D1 and D5 receptor antagonist (from here in referred to as D1-like receptors) and sulpiride, a selective D2, D3 and D4 receptor antagonist (from here in referred to as D2-like receptors) (Neve, 2013). We demonstrated a significant role for D1-like receptors in strategy formation in the FMP Y-maze, with an associated impact on working memory and cognitive flexibility. However, in the absence of reward or motivational learning, no such role was identified for D2-like receptors in exploring the maze (Cleal et al., 2020). Consequently, in models with deficits in working memory or behavioural plasticity in the FMP Y-maze, we hypothesised that increasing D1-like receptor activation would enhance these cognitive domains.

Zebrafish (*Danio rerio*) have recently emerged as a promising model of cognitive aging (Gerhard, 2007; Yu et al., 2006). Zebrafish live on average for ~3 years, but can have a life span as long as 4-5 years under laboratory conditions (Kishi, 2011). Changes in cognition are evident from ~2 years, increasing the accessibility of researching gradual senescence in a model of old age (Arslan-Ergul et al., 2016; Gerhard et al., 2002; Tsai et al., 2007; Yu et al., 2006). In humans, cellular senescence can be detected by a number of age-associated phenotypes using biological and biochemical markers. One such marker is beta-galactosidase, which in humans is only expressed in senescent cells (Dimri et al., 1995). Like humans, zebrafish also demonstrate senescence associated beta-galactosidase activity, with only faint signals detectable at 9-months old, increasing significantly once fish reach 17-months and older (Kishi et al., 2003). Additionally, like humans and rodents, zebrafish also show age-associated accumulation of oxidative proteins in skeletal muscles. Kishi, et al, showed higher levels of oxidised proteins in aged fish, at 24-months old compared to younger 3 and 6-month old fish (Kishi, 2004; Kishi et al., 2003; Tsai et al., 2007). We therefore selected 6 and 24-month old zebrafish as these developmental stages show clear differences in biological markers of aging, both in zebrafish and in humans. Also, like humans and rodents, zebrafish have homologues of neurotransmitters, associated systems and brain regions necessary to assess learning, memory and executive functions such as attention and cognitive flexibility (Horzmann & Freeman, 2016; Parker et al., 2013; Tufi et al., 2016). Ease of pharmacological manipulation and high throughput behavioural testing additionally add to the convenience and suitability of zebrafish to model the effects of aging and age-related diseases (Gerhard, 2003; Gerhard & Cheng, 2002; Keller & Murtha, 2004; Kishi, 2004).

The FMP Y-maze has been developed to assess cognition, specifically, exploration, working memory and behavioural flexibility, in a range or organisms, including zebrafish, flies, rodents and a virtual task for assessing humans (Cleal et al., 2020; Cleal & Parker, 2018; Fontana, Cleal, & Parker, 2019; Fontana, Cleal, Clay, et al., 2019). Analysis of exploration patterns has led to the observation that vertebrate species are biased towards strategies utilizing alternating left and right turns, known as an ‘alternation’ (left, right, left, right-LRLR, right, left, right, left-RLRL). Pharmacological blockade of memory forming pathways in zebrafish, revealed a role for memory processing and behavioural flexibility in the formulation of search patterns and updating exploration strategies during task progression (Cleal et al., 2020). In the FMP Y-maze, working memory is based on the ability to recall previous arm entries, similar to the alternation task of the T-or Y-maze (Stewart et al., 2011; van der Staay et al., 2011). Recording percentage-use of reoccurring patterns of four consecutive turn choices, known as tetragrams, over one hour of exploration gives rise to a ‘global’ search strategy. Similar exploration strategies, predominantly reliant on alternations, have been reported in rodents, both in the FMP Y-maze and T-maze using the same analysis (Cleal et al., 2020; Gross et al., 2011). Behavioural assessment of one hour of free exploration in the FMP Y-maze enables the identification of working memory abilities and cognitive flexibility, presenting the FMP Y-maze as a useful tool for assessing memory processing in animal models and potentially patients with disorders characterised by cognitive decline.

We used the FMP Y-maze and pharmacological agonism of D1-like receptors, to understand the role of dopamine in healthy aging in zebrafish. We compared behavioural phenotypes of 6-month old adults with aged, 24-month old adults in the early stages of senescence. Our hypothesis was that 24-month old zebrafish may have begun experiencing mild cognitive decline with the increase of age-associated biological markers. We examined changes in working memory and cognitive flexibility in a recently developed behavioural paradigm that can be used to assess both cognitive functions simultaneously with minimal handling and experimenter interference. We furthered our investigation by comparing changes in working memory of young and aging zebrafish to changes in working memory between young and old, healthy humans. Our aim was to assess the translational relevance of age-related changes in memory processing in zebrafish compared to humans.

## Materials and Methods

### Ethical statement

The University of Portsmouth Animal Welfare and Ethical Review Board guidelines were followed for all experiments carried out as part of this study, and under license from the UK Home Office (Animals (Scientific Procedures) Act, 1986) [PPL: P9D87106F]. Human experiments were conducted following approval from the University of Portsmouth Science Faculty Ethics Committee (SFEC-2019-062).

### Animals and housing

71 male and female wild type (AB) zebrafish (*Danio rerio*) aged 6-months old (young adult, n=33) and 24-months old (aged, n=38) were used to assess cognitive aging as a commonly used strain of zebrafish for investigating aging (Kishi, 2004). Previous work in our lab has shown that there are no sex differences in the FMP Y-maze and therefore we did not examine the effect of sex as part of this study (Fontana, Cleal, & Parker, 2019). Zebrafish were bred in-house and raised in the University of Portsmouth Fish Facility. Fish were housed in groups of ~10-12 fish per 2.8L tank on a re-circulating system (Aquaneering Inc., San Diego, CA, USA), aquarium-water was maintained at pH 8.4 (±0.4). Previous work from our group and extensive pilot studies were used to calculate power analysis and inform sample sizes used in this study (Cleal et al., 2020; Cleal & Parker, 2018; Fontana, Cleal, & Parker, 2019; Fontana, Cleal, Clay, et al., 2019). Room and tank temperatures were maintained at 25-27°C on a 14/10-hour light/dark cycle. From 5 days post fertilisation (dpf) fish were fed on ZM fry food until adulthood when they were fed on a daily diet of live brine shrimp (maintained at the fish facility) and dried flake food (ZM Systems, UK) 3 times/day (once/day at weekends). All fish used in this study were experimentally naïve. Once behavioural testing had been completed test fish were euthanized by rapid cooling (immersion in 2°C water), followed by decapitation and excision of the brain for downstream processing.

### Drugs

To study the effect on working memory, cognitive flexibility and mRNA expression of dopamine receptors, adults were incubated for 30 min in 35 μM (10 mg/L) of the selective dopamine D1/D5 receptor partial agonist SKF-38393 hydrochloride (Medchemexpress). Due to the water solubility, drug stock solutions were made by dissolving SKF-38393 in aquarium water. Drug administration was adapted from (Naderi et al., 2016), in which individual fish were entirely immersed in drug solution or aquarium water (control) by gently netting fish from home tank to a 400 mL beaker filled with 300 mL of drug or water and covered with a lid to prevent fish from escaping. Fish were netted into the beaker 30 mins prior to behavioural testing. Treatment time was based on previous studies with dopaminergic agonists, demonstrating that 30 mins of free-swimming during drug immersion was sufficient for drug to influence zebrafish behaviour (Irons et al., 2013; Naderi et al., 2016). Immediately following treatment fish were transferred into the FMP Y-maze.

### FMP Y-maze

The protocol was carried out as described in our previous papers (Cleal, et al., in press; (Cleal & Parker, 2018; Fontana, Cleal, & Parker, 2019; Fontana, Cleal, Clay, et al., 2019). Fish were recorded in a Y-maze for 1 hour. If the fish were adopting a random search strategy, it would be predicted that the distribution of tetragrams over a 1-hour period would be approximately stochastic (i.e., the relative frequency of each tetragram would be ~6.25%). *Figure 1* shows an example of a series of movement sequences performed in the FMP Y-maze, a combination of which provide a picture of the global search strategy used. To minimise stress, fish handling and experimenter visibility were both kept to a minimum. Behavioural testing was conducted using a commercially available, fully integrated testing environment, the Zantiks AD system for adult zebrafish (Zantiks Ltd., Cambridge, UK). Tanks were black, opaque acrylic with a transparent base. A white acrylic Y-maze insert was fitted into each tank. Two Y-maze inserts could be fitted per tank. The Y-maze dimensions were as follows: L50 mm x W20 mm x D150 mm, with a 120° angle between arms. Tanks were filled with 3L of aquarium-water and placed into Zantiks behaviour units, one tank per unit. Each system was fully controlled via a web enabled device. Filming was carried out from above, which allowed live monitoring of fish within the behaviour system. Data output was automated, preventing any bias in the recording of arm entries.

**Figure 1a.**
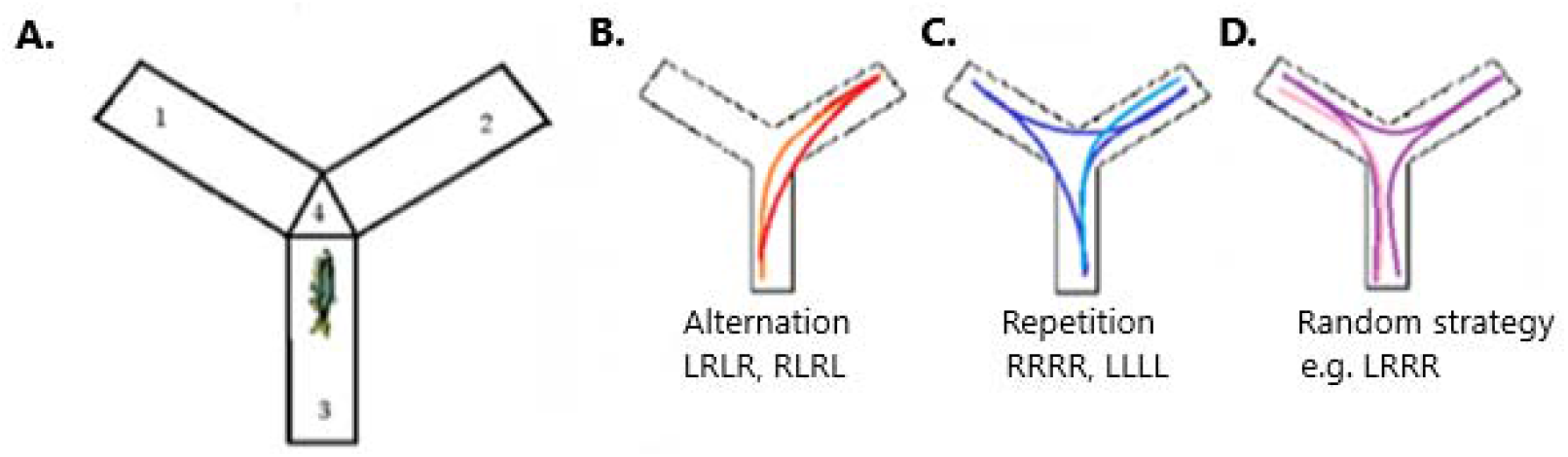
FMP Y-maze behaviour of free-swimming zebrafish. a) FMP Y-maze arena divided into three arms (1,2,3) and a neutral zone (4). b-d) Examples of movement sequences based on 16 overlapping tetragrams of left and right turns. b) Represents the dominant search strategy of alternations, made up of a series of leftright-left-right or right-left-right-left turn choices. c) Demonstrates repetitions, where an animal turns in a clockwise or anticlockwise rotation for four continuous choices, represented by RRRR or LLLL. d) An example of an alternate strategy possible using tetragram sequences. Here is shown LRRR In on anticlockwise direction, equivalent to RLLL in a clockwise direction.

**Figure 1b.**
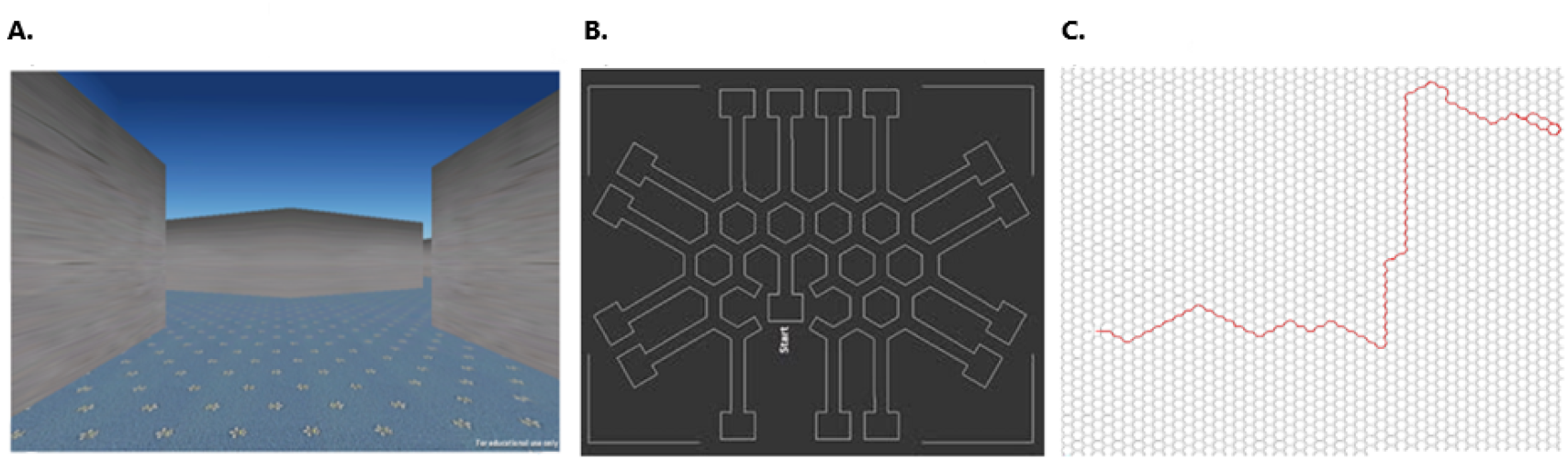
Virtual FMP Y-maze for human participants. a) In maze screenshot taken from the perspective of the participant exploring the maze. b) Schematic depicting the honeycomb pattern made up of adjoining Y-shaped mazes. c) Overlay of the left and right turns made by a participant in the maze during 5 minutes of free exploration. The left and right turns have been converted from x,y coordinates.

### Virtual FMP Y-maze

The human FMP Y-maze is based on a series of adjoining Y-shaped points, forming a honeycomb maze. At each choice point, participants must choose to turn either left or right (Figure 1A). In order to test for aging related cognitive decline, we recruited N = 80 participants, n = 40 ‘old’ (>70) and n = 40 ‘young’ (18-35). Based on power analyses from our previous studies (Cleal et al 2020), we required a minimum of n = 16 per group. An individual’s data were excluded if they completed fewer than 10 turns. Following exclusions, the final sample size for the ‘young’ group was n = 35 subjects (mean age = 24±4.79 years; 23:12 male:female), and the ‘old’ group was n = 30 subjects (mean age = 73.83±3.93 years; 18:12 male:female). The maze was uploaded onto the crowdsourcing website Prolific Academic Ltd (https://www.prolific.ac), an online platform which recruits subjects explicitly for participation in research (Palan & Schitter, 2018). Specifically, Prolific can be used to set eligibility criteria, and allows participation of eligible subjects on a first-come, first-served basis until the total number of required participants is fulfilled. We established two studies, with the eligibility criteria based on age only. Once in the maze, participants were free to explore for a total of 5 minutes, after which they were automatically logged out and, via a link, returned to the Prolific website. Turn directions of participants were reported as x, y coordinates which were converted into left and right turns and subsequently transformed into tetragram sequences. Each participant was only permitted to enter the maze once. All participants were compensated £0.65 (rate = £7.20/hour) for taking part.

### RNA Extraction and cDNA Synthesis

Immediately following behavioural testing, fish were euthanised in ice water, brains were removed, snap-frozen in liquid nitrogen and stored at −80°C until further use. RNA was isolated using the RNeasy Micro kit (Qiagen) as described in the manufacture’s protocol. Upon purification, the quality and concentration of all samples were assessed using the NanoDrop ND-1000 (Thermo Scientific). The purities of acceptable RNA samples (as measured by 260:280 and 230:260 absorbance ratios) were equal to or greater than 1.8. All samples were therefore of sufficient quality for expression-level analysis. Total cDNA was prepared using Applied Biosystems High Capacity RNA-to-cDNA Kit for RT-qPCR. Each 20 μL reaction was diluted 10-fold in nuclease-free water and used as the template for the real-time qPCR assays.

### Quantitative Real-Time PCR (RT-qPCR)

Quantitative real-time PCR (RT-qPCR) assays were used to validate relative gene expression based on SYBR green detection. Primers used in this study were predesigned primers from qPrimerDB (Lu et al., 2018) or based on previous studies (Parker et al., 2016; Tang et al., 2007). Primers were synthesised by Invitrogen and are listed in Table 1. The Roche LightCycler® 480 High Resolution Melting Master mix and the LightCycler® 96 (Roche Life Science) were used to amplify and detect the transcripts of interest. Thermal cycling conditions included an initial denaturation step at 95°C for 600s (recommended by manufacturer), 40 cycles of 95°C for 15s, 58°C for 20s and 72°C for 35s followed by melt curve analysis to confirm product specificity for all transcripts. Primers were tested with melt curve analysis and negative reverse transcription (RT) controls and negative template controls to optimise reaction conditions to generate a single melt peak in control samples, check for genomic contamination in negative RT controls and primer dimers in negative template controls. Elongation factor 1 alpha (*eelf1a*) was used as a housekeeping gene (Tang et al., 2007). Gene-expression levels were calculated using delta-delta CT method and normalised to *eelf1a*. Changes in expression were presented as means ± SD of fold change to control group (n = 4-6 per group; assayed in duplicate).

**Table 1.**
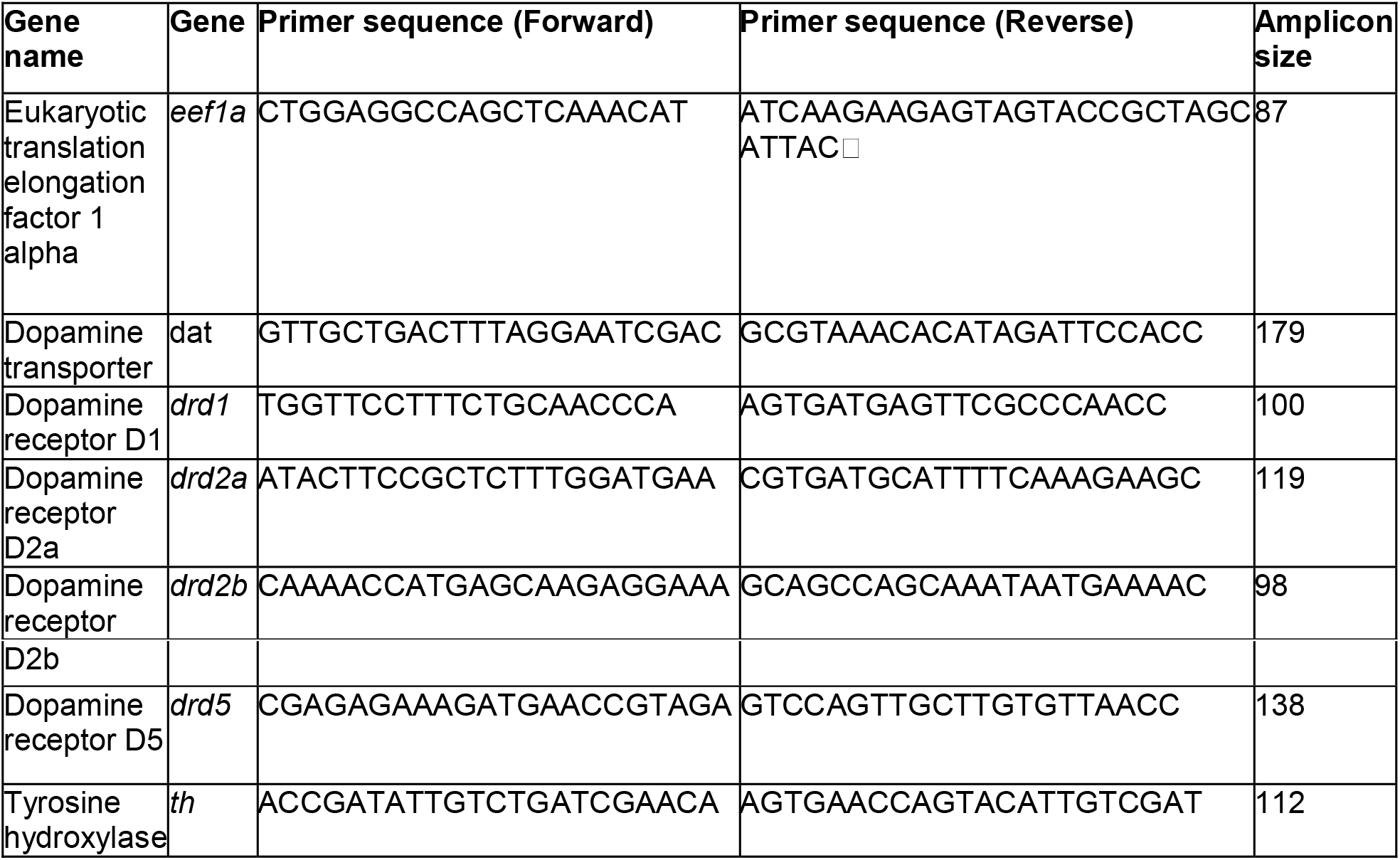
Primer sets used for qPCR

### Oxygen consumption measurements

Standard oxygen consumption rates of individual fish were measured based on wet weight (g) and amount of oxygen consumed during 1 h of free exploration of the FMP Y-maze. Method for measuring oxygen consumption were adapted from (Voutilainen et al., 2011). Briefly, each behaviour tank was filled with exactly 3 L of aquarium-water and an initial reading of oxygen saturation were recorded using a HQ30D portable dissolved oxygen meter. Zebrafish were netted directly into the maze which was covered with parafilm to create a closed system. Tanks were immediately placed into the behaviour unit and the trial started. At the end of 1 h of exploration oxygen readings were recorded for each fish. Zebrafish were then briefly anaesthetised using Aqua-Sed anaesthetic treatment (Aqua-Sed^™^, Vetark, Winchester, UK) in accordance to manufacturer guidelines. A wet weight was recorded for each fish in grams. Oxygen consumption was expressed as mg/L O_2_ x g^-1^ x h^-1^, in which g^-1^ is per gram, based on the wet weight of each fish and h^-1^ is per hour, based on 1 hour of exploration. Temperature readings were recorded pre and post behaviour to ensure that temperature did not fluctuate significantly over the course of the behavioural task, as this could impact metabolic rate. However, temperature only fluctuated by ±1°C which has been shown to be too small a change to influence oxygen metabolism (Barrionuevo & Burggren, 1999; Okomoda et al., 2020).

### Statistical analysis

The primary endpoint for analysis of exploration patterns in the FMP Y-maze was the number of choices for each of the 16 tetragrams as a proportion of total turns (percentage use). Based on previous research we were interested particularly in the proportion of choices that represented alternations (LRLR, RLRL) and repetitions (RRRR, LLLL) as these tetragrams were the most observed. All analyses were carried out using IBM SPSS statistics version 25 and GraphPad Prism version 8, with all graphical representations completed in GraphPad Prism. Analysis of paired tetragram sequences (e.g. LRLR+RLRL, LLLL+RRRR, etc) for a single age group was carried out using a One-way analysis of variance (ANOVA) followed by Tukey’s *post-hoc* test. Comparison of tetragram frequency distributions for age x treatment were analysed using Two-way ANOVA. Data were represented as mean ± standard error of the mean (SEM). For subsequent tetragram analyses, we were interested in putative changes in strategy during the search period, and we therefore included “time” as the within-subjects factor. We also included “total turns” as a covariate in all analyses, in order to control for general activity levels in statistical models. For age groups in which drug was added, we included drug (present vs absent) as a between-subjects’ factor. Data for time were represented as mean ± SEM. For all data sets the Shapiro-Wilk test of normality was conducted. Parametric and nonparametric tests were performed as appropriate for each analysis as detailed below. Comparison of percentage use of alternations and repetitions for each age and treatment group were analysed using the Kolmogorov-Smirnov test for non-normally distributed data sets and the unpaired t-test for normally distributed data sets. Data were represented as the mean ± SEM. Oxygen consumption was analysed using a One-way ANOVA for age and Two-way ANOVA followed by Sidak’s multiple comparison *post-hoc* test for age x treatment. Data were represented as mean ± SEM. Body mass was analysed using unpaired t-test. Data were represented as mean ± SEM. Locomotor data were analysed for age x treatment using Two-way ANOVA followed by Tukey’s *post-hoc* test. Data were represented as mean ± SEM. qPCR data were all tested for normality using the Shapiro-Wilk test, as above. Normally distributed data were analysed using unpaired t-test, non-normally distributed data were analysed using Mann-Whitney test. Data were represented as mean ± standard deviation (SD). Alpha values of *P* ≤ 0.05 were considered statistically significant.

## Results

### Aging zebrafish show mild cognitive decline in the FMP Y-maze

Tetragram analysis revealed that global strategy (percentage use of each tetragram over the entire trial) relied on a similar pattern of turn choice, however, key differences were observed in the use of alternations (LRLR, RLRL) and repetitions (LLLL, RRRR). *Figure 2* shows tetragram frequency distribution for 1 h of free swimming in the FMP Y-maze for 6-month old v 24-month old zebrafish. Equal distribution of all possible tetragram configurations would be characteristic of a random search strategy (100% turns completed/16 tetragram configurations = 6.25%), whereas choices made above the 6.25% threshold are considered intentional and part of a global strategy. Most tetragrams were used randomly with bars falling below 6.25% (represented by the dashed line in *Fig 2*). 6-month old fish used alternations significantly more than all other search strategies (One-way ANOVA, *F* (7, 280 = 36.85, *p* <0.0001). However, there was a significant interaction between age and turn choice (Interaction, *F* (1, 68) = 11.94, *p* = 0.0009) with Tukey’s *post hoc* test showing 6-month old zebrafish used alternations significantly more than 24-month old zebrafish (95% CI diff = 3.32-14.27, p = 0.0004***). This reduction in alternations in 24-month old zebrafish resulted in a change in the global strategy to split dominance between two tetragram sequences, alternations and repetitions, instead of just alternations as seen with the 6-month old zebrafish.

**Figure 2.**
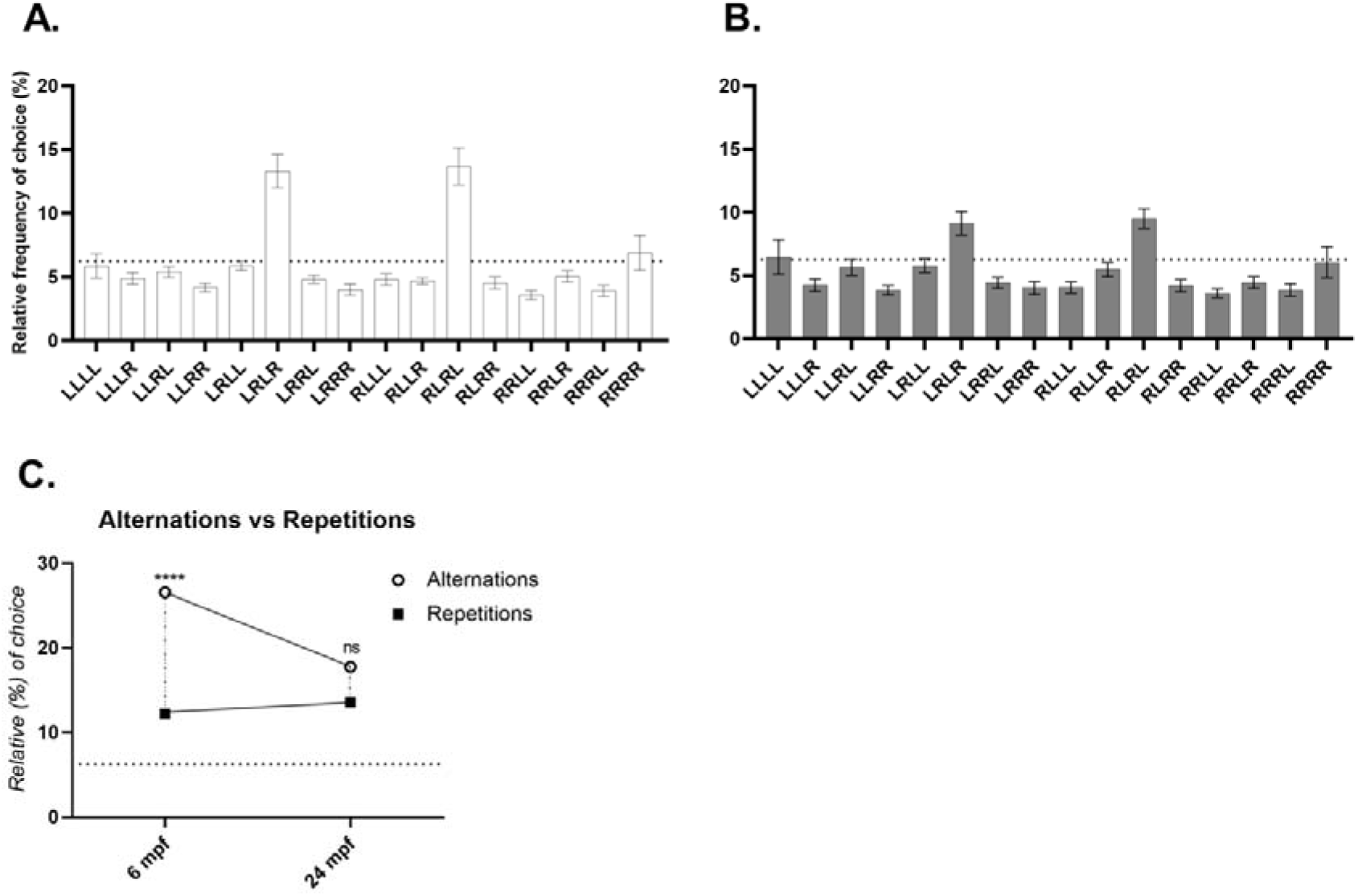
Global search strategy after 1 h of exploration in the FMP Y-maze. a) Percentage use of each tetragram sequence by 6-month old zebrafish in the FMP Y-maze (n=18), demonstrating clear dominant use of alternations. b) Percentage use of tetragram sequence by 24-month old zebrafish in the FMP Y-maze (n=20) with comparable use of alternations and repetitions. c) Alternations versus repetitions for 6-month old and 24-month old zebrafish. Normality analysed using Shapiro-Wilk test. Normally distributed data were analysed using One-way ANOVA. The dashed *line* denotes chance performance (approximately 6.25%). *** *p* ≤ 0.001, **** *p* ≤ 0.0001, ns – not significant. Error bars are mean ± SEM. Single data points are mean ± SEM.

### Treatment with D1/D5 agonist, SKF-38393, rescues working-memory deficit in aged zebrafish

Having shown a deficit in working memory of 24-month old zebrafish compared to 6-month old counterparts, shown by a change in global exploration strategy, we pretreated zebrafish with D1/D5 agonist SKF-38393 for 30 mins prior to testing in the FMP Y-maze. SKF-38393 treatment had no effect on global strategy in 6-month old zebrafish, with both alternations and repetitions showing no effect between treated (n=15) and untreated (n=18) fish (Two-way ANOVA, *F* (1, 496) = 0.44, *p* = 0.5078). However, treatment of 24-month old zebrafish caused a significant increase in the use of alterations (*KS, D* = 0.33, *p* = 0.0297), without affecting use of repetitions (*KS, D* = 0.14, *p* = 0.8582) (*Figure 3*). Working memory, as measured by global strategy showed a significant effect of age (F (1,69) = 9.037, *p* = 0.037), but no significant interaction (F (1,69) = 2.546, *p* = 0.115) and no main effect of treatment (F (1,69) = 1.643, *p* = 0.204). However, Tukey’s *post hoc* tests revealed that treatment with SKF-38393 rescued the deficit in alternations between aged 24-month old and young 6-month old zebrafish (95% CI = −3.74-10.52, *p* = 0.5969) compared to controls (95% CI = 1.91-16.18, *p* =0.0073).

**Figure 3.**
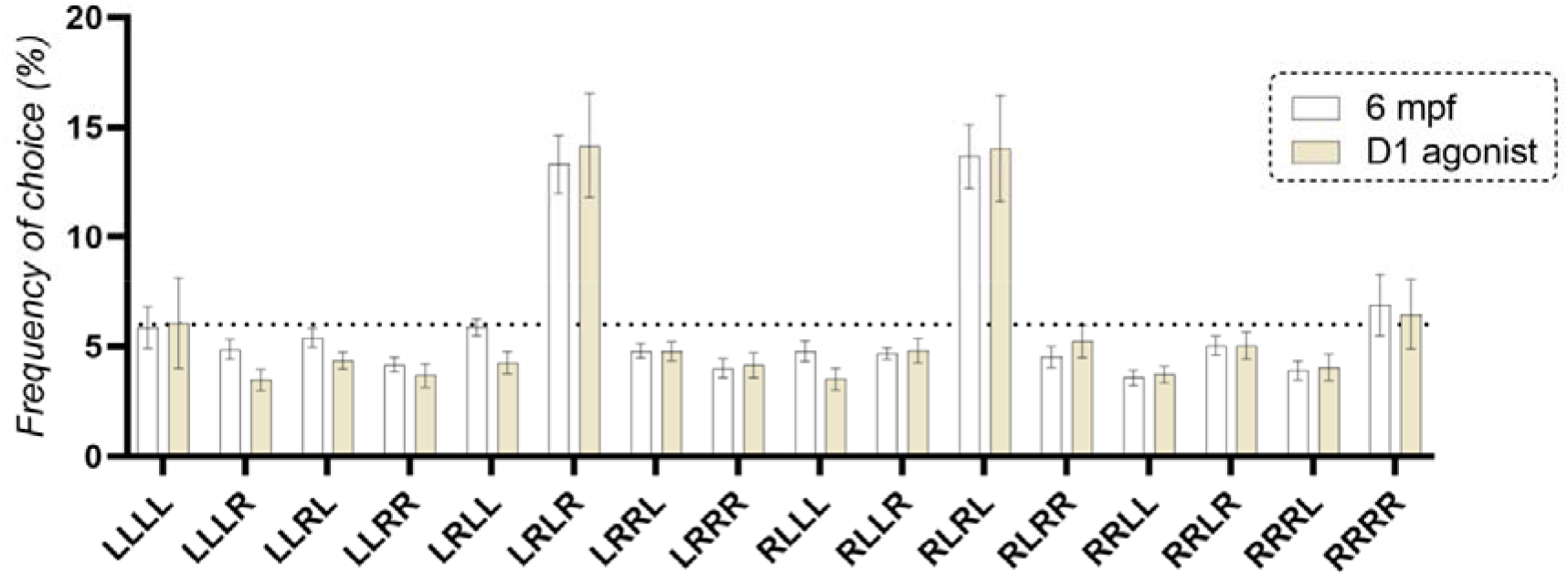
Effective pre-treatment with D1/D5 agonist SKF-38393 on global search strategy of zebrafish in the FMP Y-maze. Effect of SKF-38393 on search strategy of a) 6-month old treated compared to control zebrafish and b) 24-month old treated compared to control zebrafish after 1 h of exploration. c) Comparison of alternations and repetitions in 6-month old and 24-month old control v treated zebrafish. C – control, T – treated with D1/D5 agonist SKF-38393. Normality analysed using Shapiro-Wilk test. Normally distributed data analysed using two-way ANOVA and t-test. Non-normally distributed data analysed using Kolmogorov-Smirnov (KS) test. * *p* ≤ 0.05, ns – not significant. The dashed *line* denotes chance performance (6.25%). Error bars are mean ± SEM.

### Healthy aging impacts cognitive flexibility which cannot be recovered by treatment with SKF-38393

The FMP Y-maze is a dual-action behavioural task which enables assessment of working memory based on global strategy, but also cognitive flexibility by analysing ‘immediate’ strategies consisting of exploration patterns for the total trial divided into equal length time bins (six 10 min time bins). This enables the identification of changes in strategy over time. *Figure 4* illustrates the percentage use of each tetragram sequence per 10 mins of exploration clearly denoting differences in the use of alternations over successive 10 min exploration intervals for 6-month old zebrafish, but a diminished effect of time on alternations in the 24-month old zebrafish. As alternations have been revealed as the dominant strategy and prone to change with age and treatment, we further explored the effect of time on alternation use. 6-month old control zebrafish showed the greatest effect of time on alternations as demonstrated in *Figure 5*. From the initial 10 mins of exploration, alternations were already used above random selection and continue to rise significantly with each successive time bin (One-way ANOVA, *F* (5, 210) = 4.33, *p* = 0.009). The maximum mean difference between time bins was 9.7%, indicating that the alternation strategy was not static, but was altered in response to a constant environment. 24-month old aging zebrafish demonstrate a deficit in the ability to adapt their strategy over time, as shown by the stable use of alternations in consecutive time bins (One-way ANOVA, *F* (5, 234) = 1.35, *p* = 0.2449), with a maximum mean difference of 3.6% between time bins. Treatment of 6-month old zebrafish with SKF-38393 did not affect global strategy as seen in *Figure 3*, however, it did appear to have a dampening effect of changes in alternation use over time (*Figure 5*). Though still significant, the effect was greatly reduced compared to controls (One-way ANOVA, *F* (5, 164) = 2.73, *p* = 0.0215). In 24-month old zebrafish treated with SKF-38393 the global use of alternations was increased to a performance level equivalent to 6-month old controls; however, the drug treatment was unable to restore adaptability of search over time, resulting in a stable strategy (One-way ANOVA, *F* (5, 208) = 0.185, *p* = 0.9681), which had a similar maximum mean difference of 2.9% between time bins.

**Figure 4.**
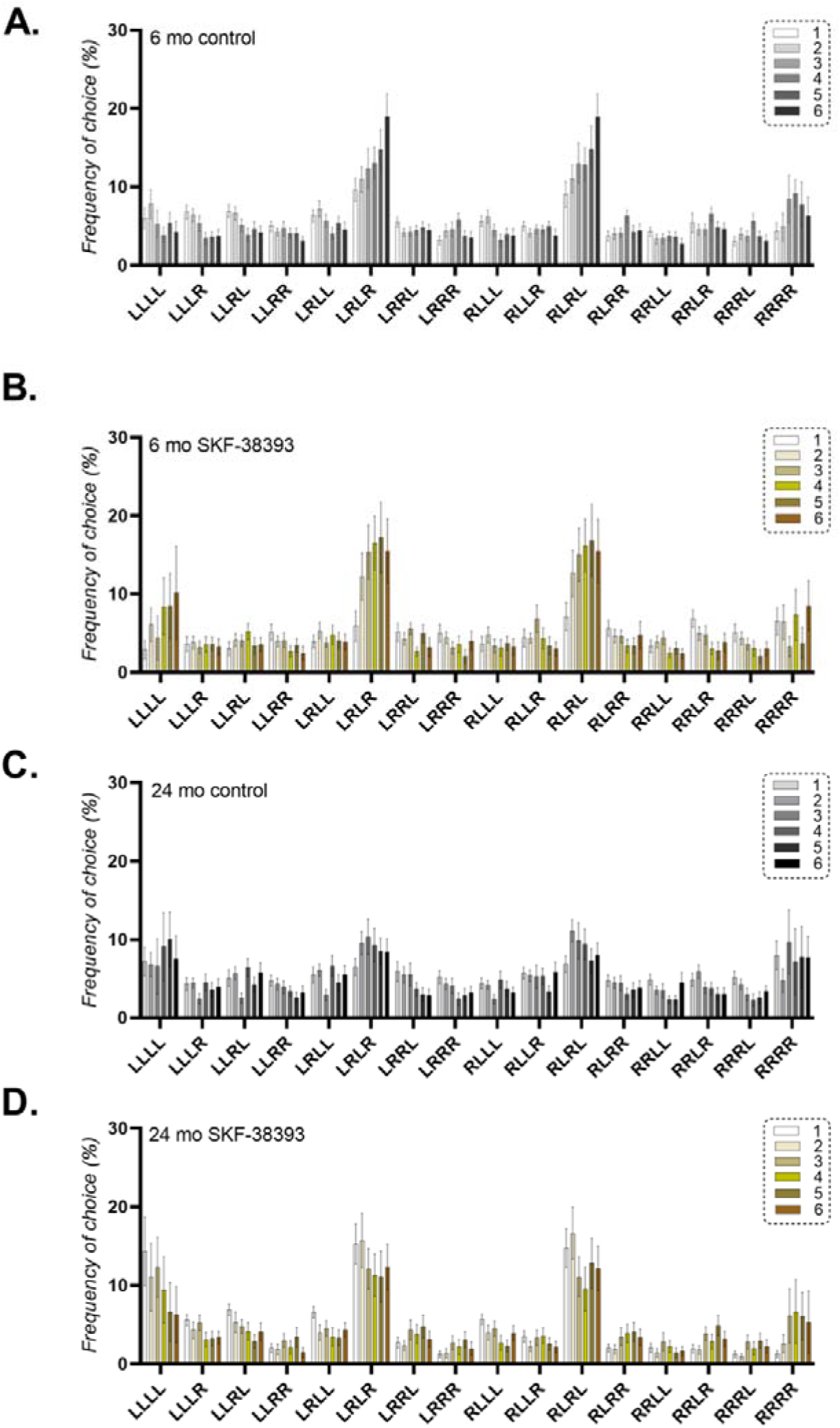
Shows the use of each tetragram per 10 min time bin for a 1 h trial of free FMP Y-maze exploration. a) Depicts tetragram use for 6-month old adult controls compared to b) SKF-38393 treated. Both groups clearly illustrate an increasing use of alterations across successive time bins. c) Shows the same frequency distribution for 24-month old aging adult controls versus d) SKF-38393 treated. Although the agonist treated group shows an increased percentage of alternations per time bin compared to controls, both groups have a heavily blunted effect of time, resulting in almost equal use of alternations in each 10 min time bin. Error bars are mean ± SEM.

**Figure 5.**
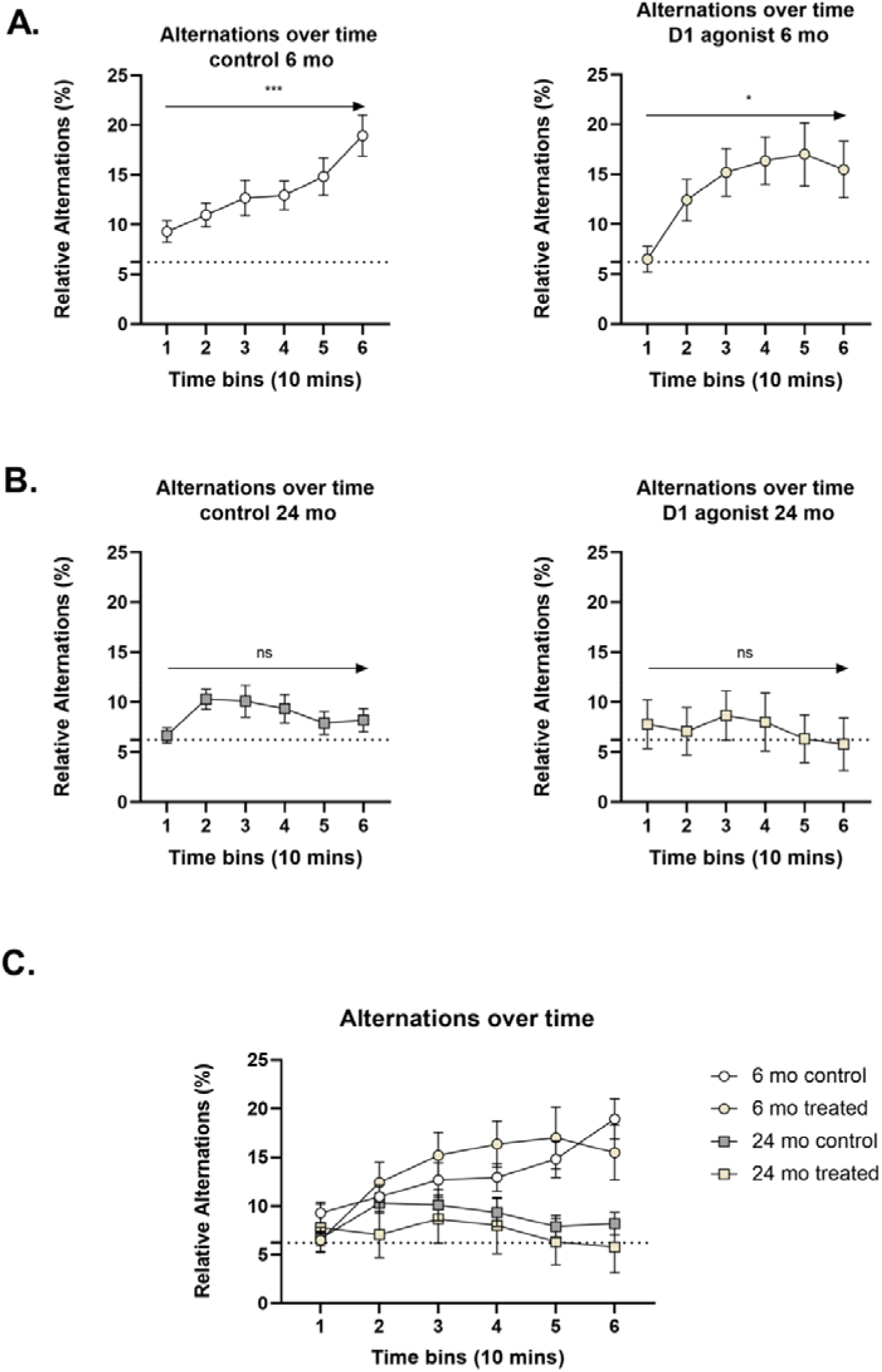
Percentage use of alternations in successive 10 min time bins of exploration a) 6-month old controls (left) and treated with SKF-38393 (right) and b) 24-month old controls (left) and treated (right). c) Combined effect of time on percentage use of alternations for all ages and treatment groups. Analysed using one-way ANOVA. * *p* ≤ 0.05, ** *p* ≤ 0.01, ns – not significant. The dashed *line* denotes chance performance (6.25%). Error bars are mean ± SEM.

### Metabolic rate is not a factor in changing search strategy with age

There appeared to be a substantial size difference between age groups, therefore, to identify if metabolism played any role in the change in search strategy, wet weight and oxygen consumption over the course of the trial were recorded as an indirect measure of metabolism (Nelson, 2016). We identified a significant difference in wet body mass (*t* = 7.195, df = 19, *p* <0.0001) with a mean wet weight of 0.39g for 6-month old and 0.83g for 24-month old zebrafish. We furthered this analysis by comparing oxygen consumption between age groups and found no significant difference (*t* = 1.660, df = 8, *p* =0.1356). However, there was an effect of treatment on oxygen consumption (Two-way ANOVA, *F*(1, 17) = 11.41, *p* = 0.0036), Sidak’s multiple comparison *post-hoc* test revealed that treatment with D1/D5 agonist SKF-38393 caused a significant decrease in oxygen consumption in 6-month old (95% CI = 0.433-1.787, *p* = 0.0012), but had no effect on 24-month old fish (95% CI = −0.631-0.535, *p* = 0.9952) (*Figure 6*).

**Figure 6.**
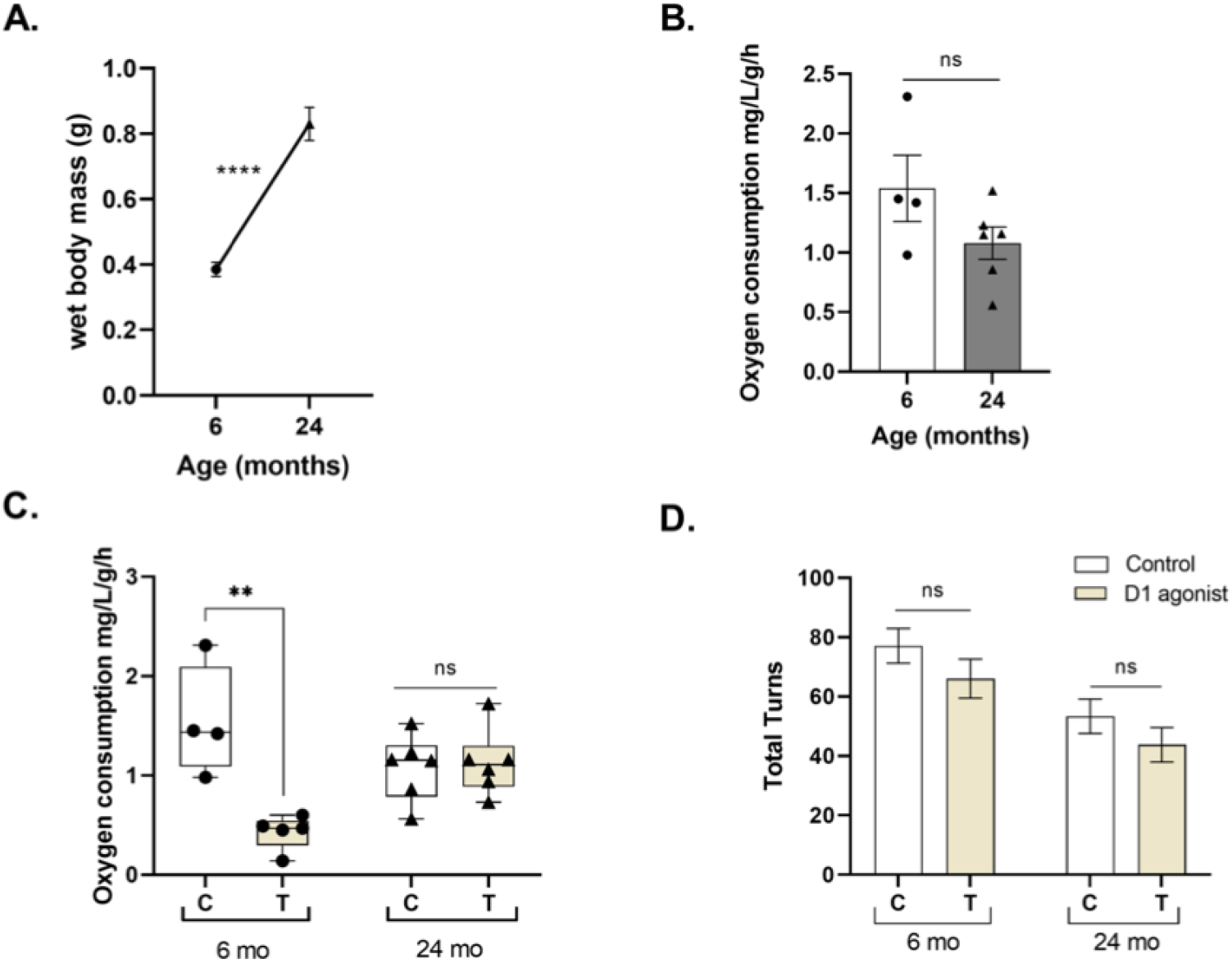
a) The mean difference in body mass between 6-month old (n=10) and 24-month old (n=12) zebrafish. b) Locomotor activity measured by total turns completed in 1 h of exploration. c) Oxygen consumption of 6-month old controls (n=4) compared to 24-month old controls (n=6). d) Effect of SKF-38393 on oxygen consumption over 1 h of exploration in 6-month old and 24-month old control v treated. Normality analysed using Shapiro-Wilk test, normally distributed data analysed by Two-way ANOVA (b,d) or unpaired t-test (a,c). Data for a and b are mean ± SEM, data for c and d are mean ± SEM; * *p* ≤ 0.05, ** *p* ≤ 0.01, **** *p* ≤ 0.0001, ns – not significant.

### Regulation of dopaminergic gene expression by aging

We investigated the effect of age on the dopaminergic system by analysing expression changes between 6 and 24-month old zebrafish. *Figure 7* shows the qPCR data of relative gene expression from whole brain tissue. We found no significant difference between 6 and 24-month old zebrafish for *dat* (*U* = 4, *p* =0.1667), *drd1* (*t* = 1.25, df = 8, *p* =0.2460), *drd2a* (*t* = 1.20, df = 7, *p* =0.2699), *drd2b* (*t* = 0.21, df = 7, *p* =0.8434) *or th* (*t* = 0.55, df = 7, *p* =0.5985) mRNA expression levels. However, there was a significant effect of age on *drd5* (*t* = 2.87, df = 7, *p* = 0.0239) expression.

**Figure 7.**
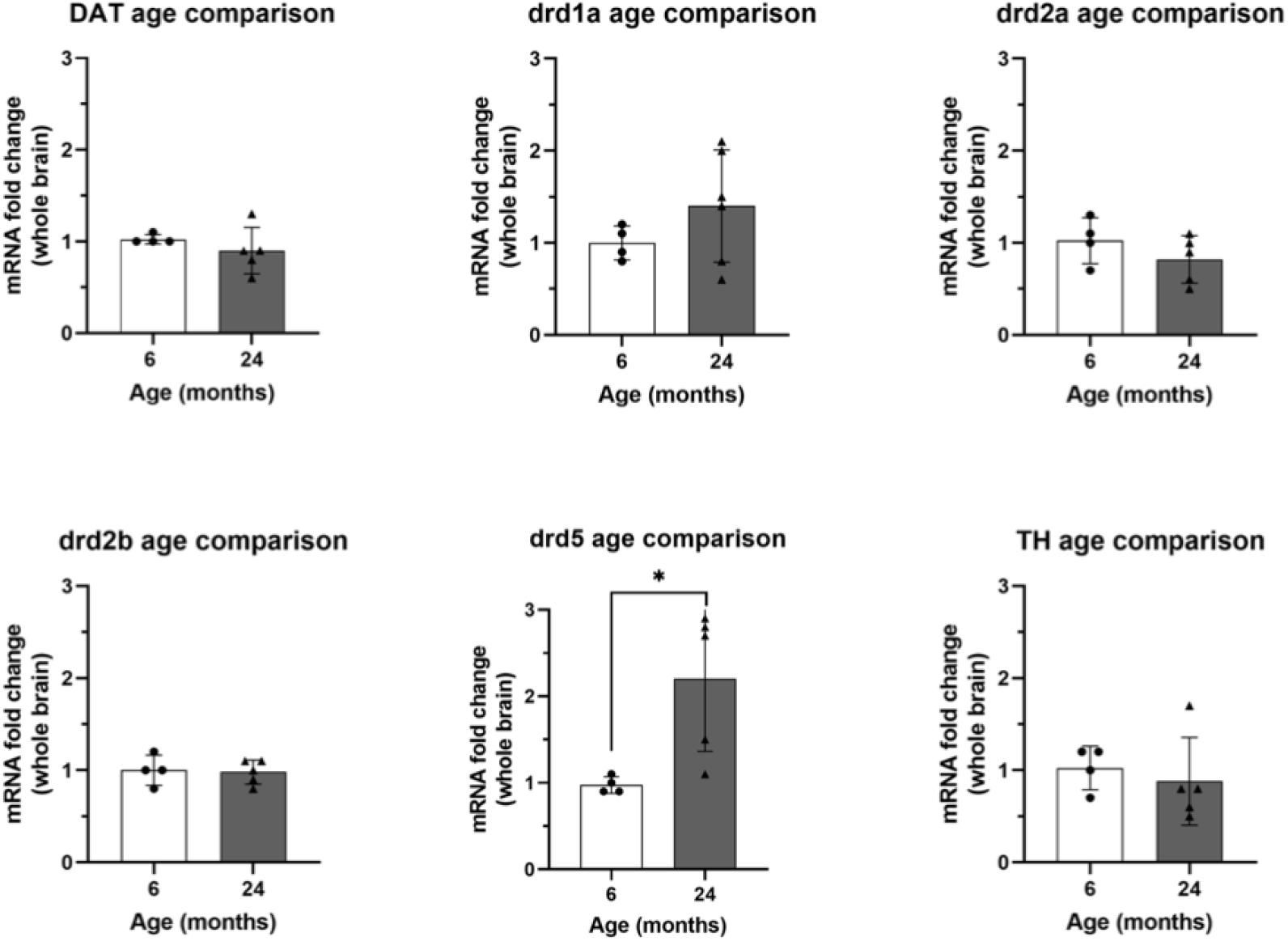
Quantitative real-time PCR analysis showing variations in the relative amounts of *dat, drd1, drd2a, drd2b, drd5* and *th* mRNA in whole brain tissue extracted from 6-month old controls (n=4) and 24-month old controls (n=6). Data were normalised to housekeeping gene *eelf1α* and defined as fold change relative to 6-month old controls. Normality was analysed using Shapiro-Wilk test. Normally distributed data were analysed using unpaired t-test, non-normally distributed data were analysed using mann-whitney test (U). All data are mean ± SD; * *p* ≤ 0.05.

### Effect of SKF-38393 on dopaminergic gene expression

The qPCR data in *Figure 8* shows that 35μM SKF-38393 was able to decrease the expression of *dat* in 6-month old zebrafish (*U* = 0, *p* = 0.0048) but not 24-month olds (*U* = 9, *p* = 0.1645). However, 30 min treatment with SKF-38393 did not elicit expression changes in *drd1* (6-months old; *t* = 1.36, df = 7, *p* =0.2170, 24-months old; *t* = 0.83, df = 10, *p* =0.4266), *drd2a* (6-month old; *t* = 1.33, df = 7, *p* =0.2259, 24-month old; *t* = 1.31, df = 8, *p* =0.2265), *drd2b* (6-months old; *t* = 1.17, df = 8, *p* =0.2753, 24-months old; *t* = 0.18, df = 8, *p* =0.8651), *drd5* (6-months old; *t* = 1.38, df = 8, *p* = 0.2064, 24-months old; *t* = 1.62, df = 9, *p* = 0.1405) or *th* (6-months old; *t* = 0.72, df = 8, *p* =0.4950, 24-months old; *t* = 0.14, df = 8, *p* =0.8894) in either age group.

**Figure 8.**
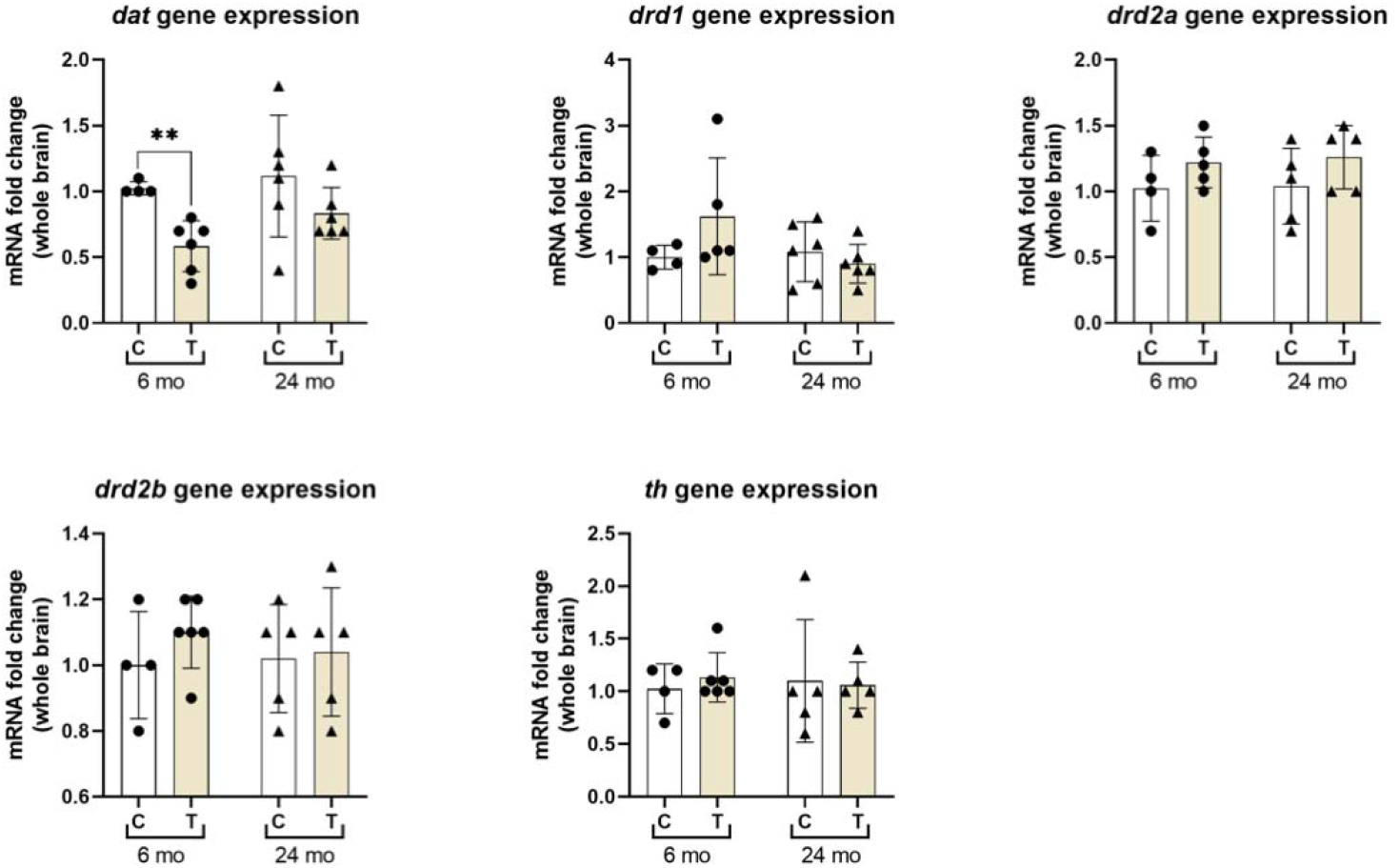
Quantitative real-time PCR analysis showing variations in the relative amounts of *dat, drd1, drd2a, drd2b, drd5* and *th* mRNA in whole brain tissue of controls and treated groups exposed to 35 μM SKF-38393 in 6-month old controls (n=4) and treated (n=5) and 24-month old controls (n=6) and treated (n=6) zebrafish. Data were normalised to housekeeping gene *eelf1α* and defined as fold change relative to controls. Normality was analysed using Shapiro-Wilk test. Normally distributed data were analysed using unpaired t-test, non-normally distributed data were analysed using mann-whitney test (U). All data are mean ± SD; ** *p* ≤ 0.01.

### Healthy aging humans show mild cognitive decline

Having shown a deficit in working memory of old versus young zebrafish, to understand the translational relevance of these findings, we assessed working memory in healthy populations of young and old humans in a previously developed virtual FMP Y-maze, which is analogous to the animal version (Cleal, et al., 2020). To establish clinically relevant findings, we ran participants in the FMP Y-maze for 5 minutes of free exploration. Like zebrafish, humans showed dominant use of the alternation strategy and similarly there appeared to be a significant reduction in the use of alternations in older (70+ year-olds) compared to younger (18-35 year-old) adults (*F* (1, 65) = 7.175, *p* = 0.009) (Figure 9). Thus, we demonstrated a deficit in working memory in healthy aged adults compared to their younger counterparts.

**Figure 9.**
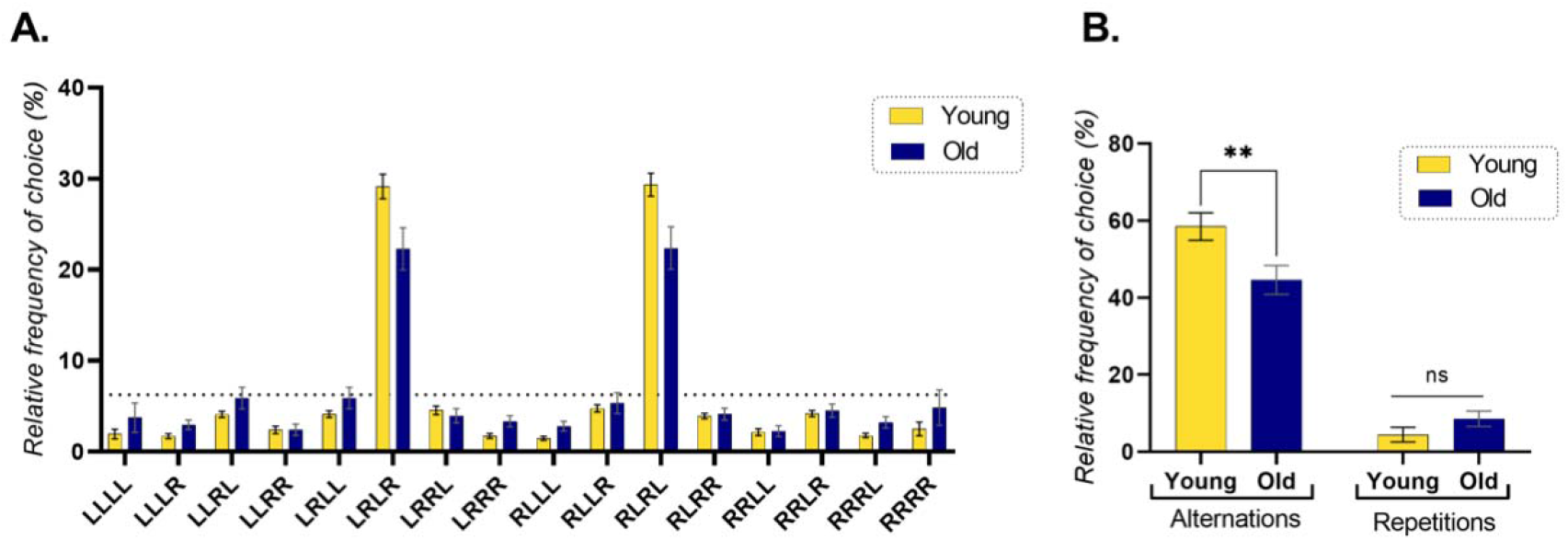
Global search strategy after 5 minutes of exploration in the human virtual FMP Y-maze. a) Percentage use of each tetragram sequence by young adults aged 18-35 years old (n=35) compared to older adults aged 70+ years old (n=32). Both demonstrating dominant use of the alternation strategy. b) Comparison of young and old aged groups and the use of total alternations (LRLR+RLRL) and total repetitions (LLLL+RRRR). Analysed using LMM. The dashed *line* denotes random choice selection at 6.25%. Error bars are mean ± SEM. ** *p* < 0.01.

## Discussion

Using a zebrafish model of aging we have investigated changes in working memory and cognitive flexibility between young adulthood and aging, suggesting mild cognitive decline. The present findings indicate that working memory and strategy changes are both impaired in aged zebrafish; however, acute exposure to SKF-38393, a selective dopamine D1/D5 receptor partial agonist, enhanced working memory in aging, but had no detectable effect on behavioural flexibility. Real-time qPCR analysis identified an up regulation of *drd5* receptor in 24-month old zebrafish compared to 6-month old counterparts. Treatment with SKF-38393 caused a down regulation of *dat* mRNA expression in 6-month old adults, but no effect on aged adults, supporting the role of *dat* in regulating dopamine availability and cognitive flexibility. These findings provide characterisation of cognitive changes in healthy aging which can be partially rescued by activating D1-like receptors, promoting the role of maintaining working memory, but not behavioural flexibility. This study, to our knowledge, provides the first examination of changes in dopaminergic gene expression in the aging zebrafish brain. To address the translational relevance of the zebrafish model of aging, we examined working memory of healthy adults using an analogous version of the FMP Y-maze adapted for humans. We identified a similar reduction in alternations in old versus young adults, comparable with the changes seen in zebrafish. Thus, our findings demonstrate for the first time that zebrafish faithfully replicate aspects of healthy aging seen in humans, that results in a decline in working memory with old age.

### Changes in working memory in healthy aging zebrafish

Younger adult zebrafish have demonstrated a very specific global strategy used to explore the FMP Y-maze. Using tetragram sequence analysis, we identified patterns of choice selection of left and right turns in 6-month old adults that use alternation sequences (LRLR, RLRL) for more than 25% of the global search strategy, four times the use of random selection. All other sequences were used at a level equivalent to chance selection. Prior wsork from our group has demonstrated that changes in global spatial activity patterns, particularly those relating to alternations and repetitions, are representative of changes in working memory processing (Cleal, et al., 2020). Previous work has also demonstrated similar patterns of alternations in young adult zebrafish ranging from 3-6-months old (Cleal & Parker, 2018; Fontana, Cleal, & Parker, 2019; Fontana, Cleal, Clay, et al., 2019). Aging zebrafish at 24-months old have, however, demonstrated a marked change in global spatial activity patterns reducing the use of alternations by ~8%, bringing the use of alternations and repetitions almost in line at 18% and 16% respectively. This deficit in alternations may represent an inability to recall which arms of the maze have previously been entered and/or the order of entry, a process which has been shown to be dependent on working memory (Lalonde et al., 1986; Moran et al., 1995; Myhrer, 2003). These findings further support that zebrafish, like humans, have a natural decline in cognitive abilities as part of healthy aging, resulting in deficits in working memory (Berry et al., 2016; Dreher et al., 2008; Goldberg, 2017; Salthouse, 2009).

The role of aging in cognitive decline has been well documented in humans, with many animal models replicating similar deficits in cognitive performance (Gerhard, 2007; McQuail & Nicolle, 2012). Human and animal studies, have implicated disruption of the dopaminergic system in age-related decline in executive functions such as working memory (Castner & Goldman-Rakic, 2004; Costa, 2014; Decker & McGaugh, 1991; Dreher et al., 2008; Godefroy et al., 1989). To this end we pre-treated adult and aging zebrafish with the partial dopamine D1/D5 receptor agonist to identify what role D1-like receptors play in the cognitive decline in healthy aging zebrafish. Similar to animal and human studies (Castner & Goldman-Rakic, 2004; Hemby et al., 2003; Molloy & Waddington, 1988; Wang et al., 2019), we found that treatment with a partial D1/D5 agonist enhanced working memory in aging zebrafish, resulting in a rise in the use of alternations as part of the global strategy. 24-month old, treated zebrafish increased alternations by nearly 7%, bringing alternation use to a level comparable with 6-month old zebrafish. The effect of SKF-38393 was age specific, as no such changes in working memory were observed in 6-month old zebrafish. Our findings support previous studies, implicating that boosting D1-like receptor activation is only beneficial if there is pre-existing dysregulation within the dopaminergic system (Cools & D’Esposito, 2011).

### The role of metabolism and the effect of SKF-38393 on cognitive flexibility

Fine-scale analysis revealed that not only do fish use an overall, global strategy of alternations and repetitions to explore the FMP Y-maze, but this strategy is subject to change over time. We observed a pattern of increasing use of alternations throughout the trial in young, 6-month old adults. However, this natural tendency to modify search strategy with time ceases in aged adults. Throughout the life-span of an organism there is a continuous, dynamic equilibrium between goal stabilised and destabilised behaviours, in which an individual is required to balance focus on the current task against new information altering current goal perceptions (Cools, 2016). This cognitive flexibility has been strongly associated with the dopaminergic system and working memory (Cañas et al., 2003). In the FMP Y-maze the current goal could be perceived as foraging for potential food sources, or information seeking. This behavioural paradigm works on the basis of negative feedback (a continuous lack of food or reward) to update knowledge of the environment and inform decisions to selection appropriate behaviours, in this task reflected as search strategy. In young, healthy adult zebrafish the negative feedback loop dictates an increasing use of alternations over time. However, in line with previous studies in humans (Harada et al., 2013), aged zebrafish could not utilise negative feedback to update behaviour in response to the lack of environmental change and thus relied on the ‘immediate’ strategy, that was employed within the first 10 mins of novel exploration, as the only strategy used to explore the maze. This strategy was used regardless of how familiar the environment had become or in light of a continued lack of reward or novelty whilst exploring the FMP Y-maze. Changes in cognition is a normal aspect of healthy aging and similarly to humans, zebrafish exhibit age-related deficits in cognitive abilities (Adams & Kafaligonul, 2018). It is therefore unsurprising that in aging adults there is no longer an effect of time on alternations. The ability to adapt behaviour based on new information is fundamental to healthy cognitive processing, the disruption of which is common to psychiatric illness and neurodegenerative diseases (Pittenger, 2013; Waltz, 2017). The inability to adapt behaviour in response to the environment is of critical importance in aging conditions, and thus highlights the suitability of zebrafish to aid in informing human conditions of cognitive decline in aging.

An interesting finding from this study was that treatment with SKF38393 caused a reduction in the effect of time on search strategy in younger adults. However, aged adults showed no effect of time on search strategy, regardless of whether they were treated with SKF-38393 or not. Dopamine acts as a neuromodulator, which is essential for achieving and maintaining cognitive control functions such as flexibly adjusting goal-directed behaviours (Ott & Nieder, 2019). Via D1-like receptor activation, dopamine can modulate working memory performance and sustain goal-focused behaviour by stabilising neuronal activity and generating a high signal to noise ratio, reducing the influence of interfering (off target or distracting) stimuli (Durstewitz et al., 2000). However, in order to achieve goal-orientated behaviour with flexible adaptations a balance is required between a D1 (including both D1 and D5 receptors)- and D2-(including D2, D3 and D4 receptor) dominant state, known as Dual state theory (Cools, 2016). Dual state theory implicates intermediate neural levels of dopamine, primarily acting via D1-like receptors, resulting in low firing rate (lower energy use) and increased goal-directed behaviour as the D1-dominant state. The D2-dominant state, activing via D2-like receptors, is characterised by fast firing rates (high energy use) and behavioural flexibility, e.g. set shifting or behavioural adaptations (Cools, 2016). A shift to the D1-dominant state by pharmacological intervention, for example following pre-treatment with a selective D1/D5 receptor agonist, would result in system bias in favour of goal stabilisation, which is good for achieving the current goal (e.g. searching for food), but bad for adapting behaviour in response to new information (e.g. no food and a constant, unchanging environment, as presented in the FMP Y-maze) (Cools, 2016; Yu & Yu, 2017).

We measured oxygen consumption in control and treated groups to identify if changes in metabolic activity played a role in the change in behaviour. Supporting the theory of dual state action between D1-like-goal-directed behaviour and D2-like-flexible behaviour. We found that 6-month old controls showed the greatest behavioural flexibility, but also had the highest mean oxygen consumption of 1.54 mg/L O_2_ over 1 h of exploration. Treatment with SKF-38393 caused a significant decrease in oxygen consumption and a reduction in adaptive search behaviour over time. However, no such effect was evident in aged adults which had similar search strategies over time and similar oxygen consumption in control and treated groups. Our findings support over activation of the D1-like receptor pathways, in 6-month old adults’ treated with SKF-38393, potentially biasing the dopaminergic system in favour of a goal stabilised state, which decreased the amount of flexibility and reduced energy consumption by switching to a state with a lower firing rate in dopaminergic neurons.

The lack of change in search patterns over time in aged adults and the accompanying lack of change in oxygen consumption between treated and control groups, further supports this hypothesis. Examples of the Dual state theory have been evidenced in human, primate and computational studies (Durstewitz et al., 2000; Durstewitz & Seamans, 2008; Fallon & Cools, 2014; Ott & Nieder, 2019). Here we see that increasing D1-like receptor signalling interferes with cognitive flexibility in younger adults and is unable to restore it in aged adults. Thus, biasing away from flexible behaviour appears to be a robust mechanism; however, as we have outlined above, in order to positively influence goal-directed behaviour a specific level of receptor activation is required. It is also noteworthy that these changes do not appear to be related to locomotion as there are no changes in total turns between controls and treated fish of either age group, despite changes in metabolism in 6-month old adults and changes in global strategy in 24-month old fish in response to SKF-38393 treatment. This further supports the hypothesis that changes are mechanistic and dependent on age, opposed to physical, i.e. increased number of turns resulting in an increased number of alternations, or decreased activity decreasing oxygen consumption. Here we provide the first evidence that the Dual state theory may also apply to zebrafish. However, the modulatory effect of dopamine is complex, influenced by dopamine receptor activation and age. These findings provide further understanding of the role of the dopaminergic system in age-related changes in cognition, but more work is necessary to fully elucidate the mechanisms and their potential manipulation to improve executive function in aging adults.

### Molecular changes in dopaminergic gene expression

To further understand the role of dopamine in healthy aging we investigated the expression of genes critical to the dopaminergic system including tyrosine hydroxylase (*th*), dopamine transporter (*dat*) and the dopamine receptors *drd1, drd2a, drd2b* and *drd5*. Contrary to previous studies, we did not observe any age-dependent alterations in expression levels of *th, dat, drd1* or *drd2* receptor subtypes, however, there was a significant increase in *drd5* receptor gene expression in aged compared to young adult zebrafish. One possible explanation for the lack of detectable changes in *th, dat, drd1* and *drd2* mRNA expression may be due to the assessment of whole brains. In many mammalian studies assessing the role of aging on dopamine system regulation, gene expression is often analysed from regionspecific tissues or neuronal populations, e.g. pyramidal neurons, stellate neurons, hippocampus, prefrontal cortex, ventral or dorsal regions of the striatum (Araki et al., 2007; Godefroy et al., 1989; Hemby et al., 2003). The asymmetric distribution of dopamine receptors throughout the brain strongly correlates localization with functional specificity (El-Ghundi et al., 2007). It is therefore a possibility, that regional differences in expression may counteract one another when the system is subject to subtle changes in expression. It is also possible that differences in expression are not typical between 6 and 24-month old zebrafish, as Kishi, et al, demonstrated that age-specific changes are not consistent until 31-months old. Prior to this the extent of cellular senescence is highly variable (Kishi et al., 2003). This may similarly be the case of age-related changes in dopaminergic gene expression; however, this has not been fully characterised in adult zebrafish and highlights an area in need of further investigation.

Contrary to research on *drd1 and drd2* subtypes, relatively few studies have investigated gene expression of *drd5* in aging in the healthy brain of mammals, and to our knowledge this has not previously been investigated in zebrafish. Our finding of increased *drd5* expression is somewhat at odds with the few mammalian studies of D5 receptors in aging (Hemby et al., 2003; Rothmond et al., 2012). Rothmond, et al, found that D5 receptor mRNA expression in humans was negatively correlated with age from neonate (6 weeks old) to adult (up to 49 years old), however they noted that there was no significant difference between the age groups examined, including young adults (20-25 years old) and adults (35-50 years old). Hemby, et al, also examined age-related changes in mRNA expression of dopamine receptor subtypes in humans. They found a significant age-related decline in relative abundance of D5 receptors in CA1 pyramidal neurons, but no change in the entorhinal cortex layer II stellate neurons (Hemby et al., 2003). Additionally, the author’s noted that age-related changes in D5 receptor expression were only possible due to the ability to examine specific neuronal populations. Both studies found overall negative correlations with D5 receptor mRNA expression with increasing age, however, this was over multiple age groups differing by ~49 and ~70 years respectively (Hemby et al., 2003; Rothmond et al., 2012).

Many pharmacological studies, in mammals and fish, have identified a critical role for D1-like receptors in cognitive functioning, including working memory and cognitive flexibility (El-Ghundi et al., 2007; Messias et al., 2016; Puig et al., 2014; Stuchlik et al., 2007; Wang et al., 2019; Wietzikoski et al., 2012). However, often studies refer to the effects of “D1 receptors” or “D1 receptor agonists/antagonists”, despite the lack of ligands that can specifically select between D1 and D5 receptors. Thus contributions to cognitive function are allocated to D1-like family members (El-Ghundi et al., 2007; Nichols, 2010). Few studies have specifically investigated the role of D5 receptors on cognitive functions. One study by Carr, et al, investigated the role of D5 receptors on working memory using homozygous and heterozygous D5 receptor knockout mice (Carr et al., 2017). This study demonstrated significant alterations in cognitive function as a consequence of the loss of D5 receptors. Specifically, they identified a role for D5 receptors in spatial working memory, which correlates with the wide spread expression of D5 receptors in regions associated with the function of spatial working memory (Ciliax et al., 2000; Khan et al., 2000; Luciana & Collins, 1997). Despite work in healthy subjects identifying a negative correlation between increasing age and D5 receptor expression, the opposite has been found in models of dopaminergic disruption in which several studies have reported upregulation of D5 receptor mRNA expression. Study of L-DOPA induced dyskinesia found a significant upregulation of D5 receptor expression in response to chronic L-DOPA administration (Castello et al., 2020). Whilst examination of aberrant D2 receptor overexpression, which resulted in poor prognosis in certain tumour types, reported that increased D5 receptor expression produced an anticancer response by negatively regulating D2 receptor downstream signalling (Prabhu et al., 2019). Additionally a study of dopamine receptor subtypes in the brain of suffers of Alzheimer’s disease (AD) found that D5 receptors were the least abundantly expressed receptor in healthy controls, but was the most prominent receptor subtype in the frontal cortex of patients with AD (Kumar & Patel, 2007). These studies and others have suggested increased D5 receptor expression as a consequence of an integrated stress response, a mechanism used by cells to combat stress by manipulating the expression of genes that can potentially counteract the stressor (Pakos-Zebrucka et al., 2016; Perreault et al., 2013). In the case of increased D5 receptor expression in models of aberrant dopaminergic transmission, D5 may act as a homeostatic regulator. Thus, in our zebrafish model of aging, the increase of *drd5* in 24-month old adults may be a consequence of reduced working memory function. Improvement of this deficit with a D1/D5 receptor agonist, potentially indicates a role for dysregulation of the dopaminergic system which has hence caused an increase in *drd5* expression in the aging group. Our findings are the first to examine dopamine gene expression in healthy aging and identifies a novel role for *drd5* receptors as a mechanism to counterbalance dopaminergic disruption in a zebrafish model of aging.

Large or regionally uniform changes may be detectable with whole brain gene expression analysis. To this end we examined if there were any detectable changes in expression related to fish treated with SKF-38393. We found a significant reduction in expression of *dat* in 6-month old, treated adults compared to controls, an effect that was not replicated in 24-month old, treated fish suggesting an age specific effect. *Dat*, a plasma membrane protein exclusively expressed in dopamine synthesising neurons, plays a crucial role in regulating the amplitude and duration of dopamine-mediated neurotransmission by clearing dopamine from the synaptic cleft (Bannon, 2005; Bannon et al., 2001; Mortensen & Amara, 2003). The constitutive process of transporter trafficking of dopamine allowas for rapid up-and downregulation of cell surface transporter expression and, thus, transport activity (Gulley & Zahniser, 2003). Downregulation of *dat*, thus resulting in an increase of synaptic dopamine availability by reducing reuptake, is a likely explanation for the reduction in cognitive flexibility over time in the treated 6-month old zebrafish. Studies investigating dopamine system function modulating behaviour have identified a role for presynaptic dopamine transporters (*dats*). Studies examining cognitive flexibility in patients with Parkinson’s Disease (PD) have found that patients taking dopamine-enhancing medication (e.g. dopamine receptor agonists or L-DOPA) perform poorly on reversal or reinforcement learning tasks compared to patients that are not receiving medication (Roshan Cools et al., 2001; Rutledge et al., 2009; Waltz, 2017). Additionally it was noted that learning rates that were enhanced by dopaminergic medications only impacted positive reward learning and had no effect on negative outcome learning (Rutledge et al., 2009). These findings are consistent with other studies using methylphenidate or cocaine-both substances blocking dopamine reuptake by inhibiting *dat* (Gatley et al., 1999). (Clatworthy et al., 2009) found that young healthy subject’s orally administered methylphenidate had the least cognitive flexibility in reversal learning when associated with the greatest amount of striatal dopamine release; however, spatial working memory performance was improved with increasing amounts of striatal dopamine. Similarly studies of cocaine-use found that cognitive flexibility was selectively impaired, with some studies showing that working memory remained unaffected (Colzato et al., 2009; Stalnaker et al., 2009). Combined, these experiments support the findings from this study that increased synaptic dopamine, either by reducing dopamine transporters or treatment with dopamine-enhancing drugs, can impair behavioural flexibility in healthy subjects, or patients not treated with enhancing drugs, by causing an ‘over-dose’ of dopamine in regions with optimal dopamine levels, in relation to other dopamine depleted regions resulting in distinct functional changes, i.e. no effect on working memory, but a decrease in behavioural flexibility as seen here in 6-month old treated zebrafish compared to controls. This hypothesis would also explain the lack of effect of SKF-38393 on *dat* expression in 24-month old adults, which had already shown a deficit in cognitive flexibility in the control group, therefore pre-existing dysregulation of the dopamine system as a result of aging, prevented any ‘over-dosing’ effect causing changes in *dat* expression.

### Changes in working memory in healthy aging humans

Deficits in working memory found in aging zebrafish compared to younger counterparts, support findings from human studies (Berry et al., 2016; Dreher et al., 2008; Goldberg, 2017; Salthouse, 2009). However, to fully appreciate the translational relevance of zebrafish as a model of aging and the FMP Y-maze as a sensitive measure of cognition, we used a previously developed virtual version of the FMP Y-maze with healthy human subjects to assess working memory in aging. Previous tests in the FMP Y-maze found that humans, like zebrafish, relied on alternations as the dominant strategy for exploration in the maze (Cleal, et al., 2020). However, the sensitivity of this task to subtle changes in cognitive decline have yet to be examined. We used a virtual version of the FMP Y-maze, run using a clinically relevant protocol requiring only 5 minutes of subject participation, to assess exploration strategies. We found that young adults, aged 18-35, used alternations more than adults aged 70 and above, demonstrating an age-related deficit in working memory, in line with previous studies (Glisky, 2019; Goldberg, 2017; Pliatsikas et al., 2019). Our findings in humans, replicate those seen in zebrafish, supporting their use as a model of aging and additionally demonstrating the suitability of the FMP Y-maze as a measure of cognition that can be used to assess the same neurobiological measures in humans and fish.

## Conclusion

Our study shows that mild cognitive decline is a common component of healthy aging in both humans and zebrafish. Using the FMP Y-maze, subtle changes in working memory can be detect in aged compared to young adults. Our results are consistent with previous findings that the dopamine system plays a vital role in maintaining normal cognition and consequently appropriate behaviour selection in natural, healthy aging. We found that aged adult zebrafish have impaired working memory and cognitive flexibility compared to their younger counterparts, however, treatment with a D1/D5 agonist could improve working memory performance in the FMP Y-maze. We also identified a role for ‘over-dosing’ of dopamine and regulation of *dat* expression in subjects without dopamine depletion, which consequently resulted in reduced cognitive flexibility in healthy treated adults compared to controls. Further study is required to fully elucidate the mechanisms underlying responses to dopamine-enhancing drugs, with particular focus on regional brain changes and specific behavioural impairment and enhancement. Our work further supports the use of zebrafish as a model organism for studying behavioural changes and cognitive decline in aging, with significant translation relevance to humans.

